# Reliability-guided meta-control in intuitive physics

**DOI:** 10.64898/2026.09.14.751547

**Authors:** Aria Zhang, Samuel J. Gershman, Tomer D. Ullman

## Abstract

Intuitive physical reasoning is an important part of daily life, but the computations underlying it remain debated. Some prominent accounts propose intuitive physics relies on mental simulation, with people evolving mental scenes forward through step-by-step. However, representing every intermediate mental state in detail can be computationally costly. We suggest that intuitive physical reasoning combines mental simulation with learned predictive abstractions, and that the trade-off between these two is determined by the prospective reliability of an abstraction. We implemented the abstraction part of our proposal as a recurrent neural network model that learns a predictive abstraction by directly predicting the object’s future position from a short visual history. This network bypasses intermediate states, while also estimating its own prediction error. Our network produced simplified linear trajectories that shortcut ground-truth physical trajectories (in line with human behavior), and its error estimates were associated with human pupil size during a physical prediction task. We further developed a meta-control model that used the network’s error estimate to arbitrate between abstraction (direct prediction by the network) and simulation (running a mental simulation). The meta-control model captures key patterns in human response time and accuracy, and its predicted abstraction and simulation phases corresponded to distinct eye-movement signatures, over and above an earlier blended model that uses a hand-designed abstraction. Our results suggest that efficient physical reasoning can emerge from an interaction between learned abstractions, estimates of how reliable these abstractions are, and selective engagement of detailed simulation based on reliability estimates.

**Author summary:** From balancing dishes to knowing how an ice-cream scoop will fall and splatter, people have an intuitive sense of physics that they use in daily life. It is possible that people do this by “mental simulation”, thinking through how a scene will unfold step-by-step. But this might be slow and costly. We suggest people might combine mental simulation with shortcuts learned from experience. We built a neural network that learned to (1) predict where an object would be in the future without creating every intermediate state, and (2) estimate when its own predictions were unreliable. A meta-model can use the reliability estimate to know when to use the shortcut, and when to simulate. We compared our model to people’s behavior when they performed a physical reasoning task. Our model captured key patterns in people’s response time, accuracy, and eye movements. Our results suggest that efficient physical reasoning may combine simulation, shortcuts, and mechanisms for trading those off.

## Introduction

People can rapidly predict how physical events will unfold: whether a stack of books will topple, where a falling ball will land, whether two bikes will collide, and so on. One influential account proposes that such judgments rely on mental simulation, in which an internal model of a physical scene is evolved forward through time to predict future states [1]. Evidence for this account comes from behavioral studies, computational modeling, and neural activity [2–6].

Yet, detailed mental simulation is unlikely to be the only mechanism people use. Propagating every relevant object through every intermediate state can be computationally costly, and mental simulation is constrained by cognitive resources [7, 8]. Human judgments also show systematic departures from unrestricted simulation accounts [9]. People may reduce these costs by selectively simulating only task-relevant parts of a scene [10], or reasoning over coarse approximations rather than exact object geometry [11]. They may also combine simulation with qualitatively different, less costly strategies. In earlier work, we proposed that a blended model combining partial simulation with linear projection better captures human response times and systematic errors than either pure simulation or pure abstraction alone [12]. Other work similarly proposed that people flexibly combine heuristic rules with simulation in physical stability judgments [13]. These findings suggest that physical reasoning can trade fidelity for computational efficiency when a cheaper computation is sufficient.

The trade-off between mental simulation and shortcuts or rules can be understood as a problem of resource-rational meta-control: cognitive systems should allocate computation according to the expected benefits of a strategy relative to its costs [14–17]. This view has found general support in cognitive science, and findings show people adapt their cognitive strategies and construct simplified representations to make efficient use of limited cognitive resources [18, 19]. The expected benefit of a cognitive strategy should depend on its predictive reliability, with control shifting toward strategies whose predictions are more reliable or less uncertain [20, 21]. And indeed, people rely less on a temporally abstract predictive world model when its future-state predictions become less reliable [22], and increase costly model-based control when its accuracy advantage justifies the additional computational cost [23]. Taken together, these proposals and findings suggest that efficient reasoning dynamically allocates computation according to both its cost and expected reliability.

However, these accounts generally assume that the candidate computations or representations are already available. This raises a complementary question for intuitive physics: where do efficient abstractions come from? Existing hybrid models typically specify the alternative to simulation in advance; for example, the blended model uses a predefined linear projection [12]. But predictive shortcuts could instead be acquired through experience. Neural network models show that physical expectations can emerge from visual experience [24], that predictive representations of future physical states can be learned from dynamic visual scenes [25], and that recurrent networks can discover compact cognitive strategies rather than requiring those strategies to be specified by the researcher [26]. These findings motivate the possibility that an efficient alternative to step-by-step physical simulation could itself be learned.

Here, we develop an account of intuitive physical reasoning that balances computational efficiency and accuracy by integrating learned predictive abstraction with detailed simulation through reliability-guided meta-control. We first build a recurrent-neural-network (RNN) visual model that learns both a fast prediction of an object’s future state and an estimate of its own prediction error. Given a short visual history and a requested future time, the RNN predicts the corresponding future position directly rather than autoregressively generating every intermediate state. We use the term “abstraction” to refer to this direct temporal shortcut, in contrast to step-by-step physical simulation. Further, the added future-position prediction error provides a reliability signal for the abstraction. We use this signal in a meta-control model that relies on learned abstraction when the predicted error is low, and switches to detailed physical simulation when the predicted error is high. Even before comparing this meta-control model to human behavior, we note that it improves upon previous models that blend simulation and abstraction in several ways. First, the abstraction is learned from visual experience rather than hand-coded as a particular heuristic. Second, the decision to use abstraction or switch to simulation is governed prospectively by the abstraction mechanism’s learned estimate of its reliability, thus removing the need to continuously run a partial simulation in parallel.

We test our account using human behavior and eye movements during a physical prediction task [12] in which participants observe a falling ball, imagine the ball continuing to fall as it fades from view, and judge whether the ball will hit a goal (see Materials and Methods for details). Eye movements during this task provide a process-level measure, because gaze can reveal the spatial content of internally represented trajectories [27, 28]. We therefore asked whether model-predicted simulation and abstraction phases correspond to different oculomotor signatures. We predicted more continuous smooth-pursuit movements during mental simulation, and more rapid saccadic transitions during shortcuts/abstraction. We additionally examined pupil size as a complementary measure of reasoning effort, given its established relationship with cognitive effort allocation [29, 30].

We find converging evidence for reliability-guided meta-control in intuitive physics. The RNN produces more direct trajectories than the indirect ground-truth physical trajectories, and learns structured estimates of when its predictions are likely to be less reliable. The meta-control model uses the error signal to determine when to invoke simulation, and captures key patterns in human response times, accuracy, and eye movements. In addition, the predicted error is associated with larger pupil size, consistent with the hypothesis that more cognitive effort is invoked when switching from abstraction to simulation in order to reduce future error. Relative to an earlier blended model based on hand-specified path projection [12], the learned meta-control model also shows better correspondence with human response times and eye movements. Together, these results suggest that efficient physical reasoning can emerge from the interaction of a learned abstraction, a learned estimate of its reliability, and selective engagement of detailed simulation.

## Results

### Modeling framework

To model a learned alternative to step-by-step physical simulation, we developed an RNN that receives a short visual history and a requested future time offset, and predicts both the ball’s future position and the magnitude of the corresponding prediction error (Fig. 1; see Materials and Methods for details). Given an encoded visual history, each future position and corresponding error estimate is predicted independently at the requested offset, rather than autoregressively from earlier predictions; predicting a distant future state does not require first predicting every intermediate state. We therefore use the RNN as an abstraction mechanism that maps directly from the observed scene history to a ball’s future position.

**Fig 1.**
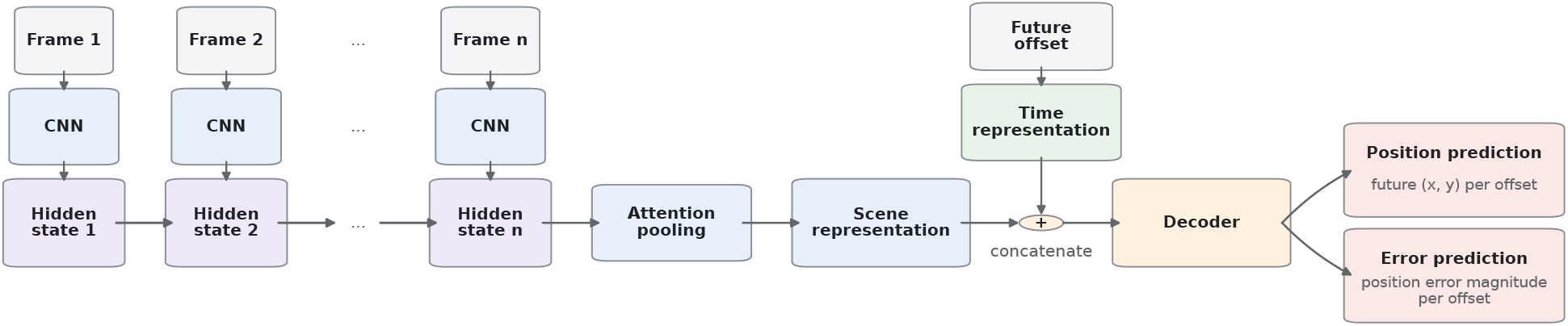
RNN model architecture. The RNN we implemented uses a short visual history to build a recurrent spatial representation, then makes time-conditioned predictions of the ball’s future position and the magnitude of its position-prediction error. Given an encoded visual history, each future position and corresponding error estimate is predicted independently at the requested offset, rather than autoregressively from earlier predictions.

The RNN-predicted error is then used to regulate abstraction and simulation in a meta-control model (Fig. 2; see Materials and Methods for details). The model uses the RNN as a fast abstraction mechanism and Pymunk as a physics simulator. Starting from the initial visual history, the controller uses the RNN to predict a distant abstraction endpoint and checks whether its predicted error remains below a threshold. If so, it accepts the reliable abstraction step and attempts to predict farther ahead. When the predicted error exceeds the threshold, the controller switches to Pymunk simulation until the resulting frames provide a new history from which abstraction is again sufficiently reliable, at which point abstraction resumes. Thus, the model uses the abstraction mechanism’s own learned error estimate as a prospective reliability signal for deciding when to trust abstraction and when to engage physical simulation. An example of the RNN’s successive future predictions and the resulting meta-control trajectory is shown in Fig. 3.

**Fig 2.**
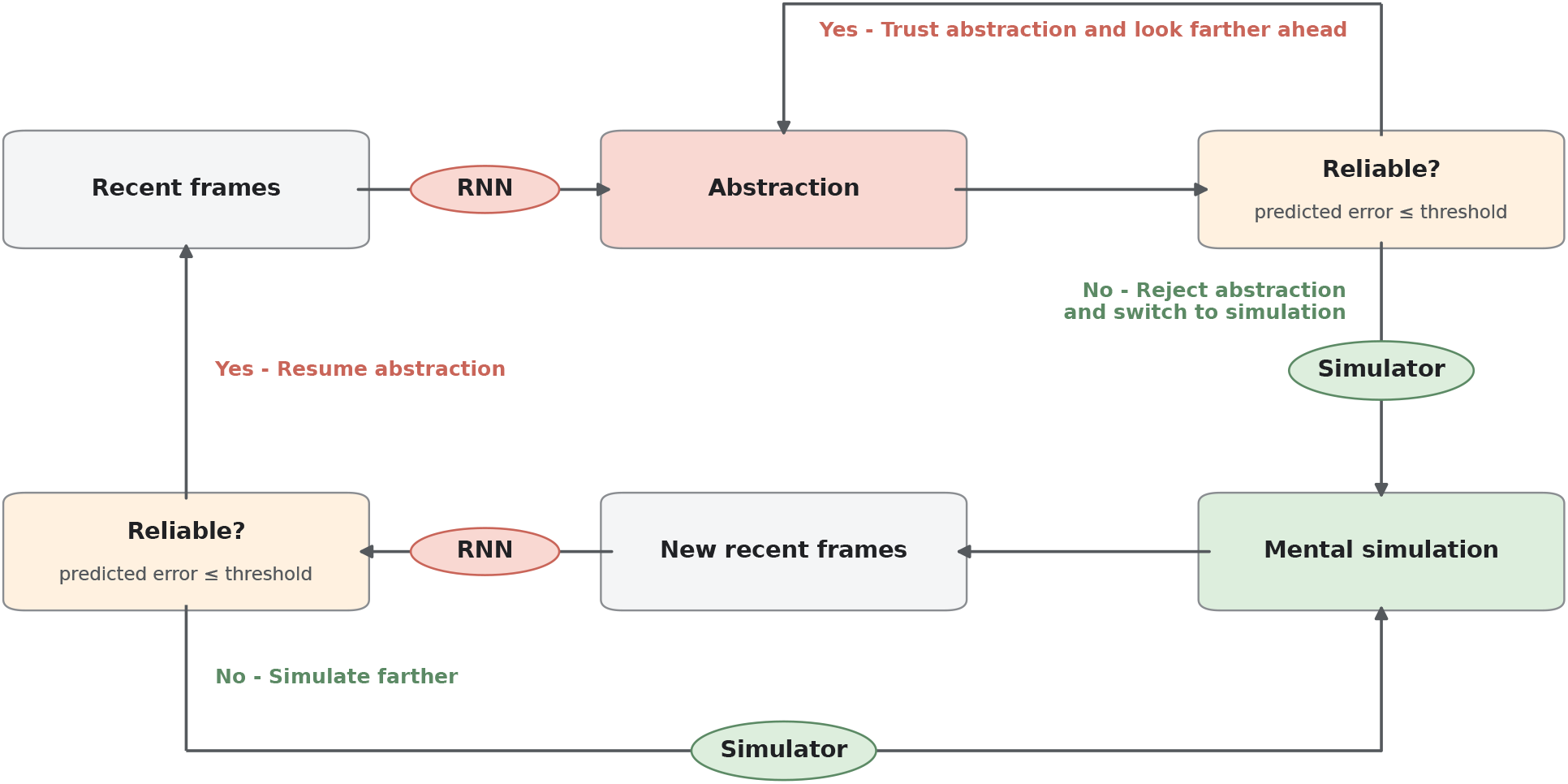
Meta-control model. Starting from the initial visual history, the controller accepts increasingly distant RNN-predicted abstraction endpoints while their error estimates remain below a threshold. When an error estimate exceeds the threshold, the controller switches to simulation until the simulated frames provide a new history from which abstraction is again sufficiently reliable.

**Fig 3.**
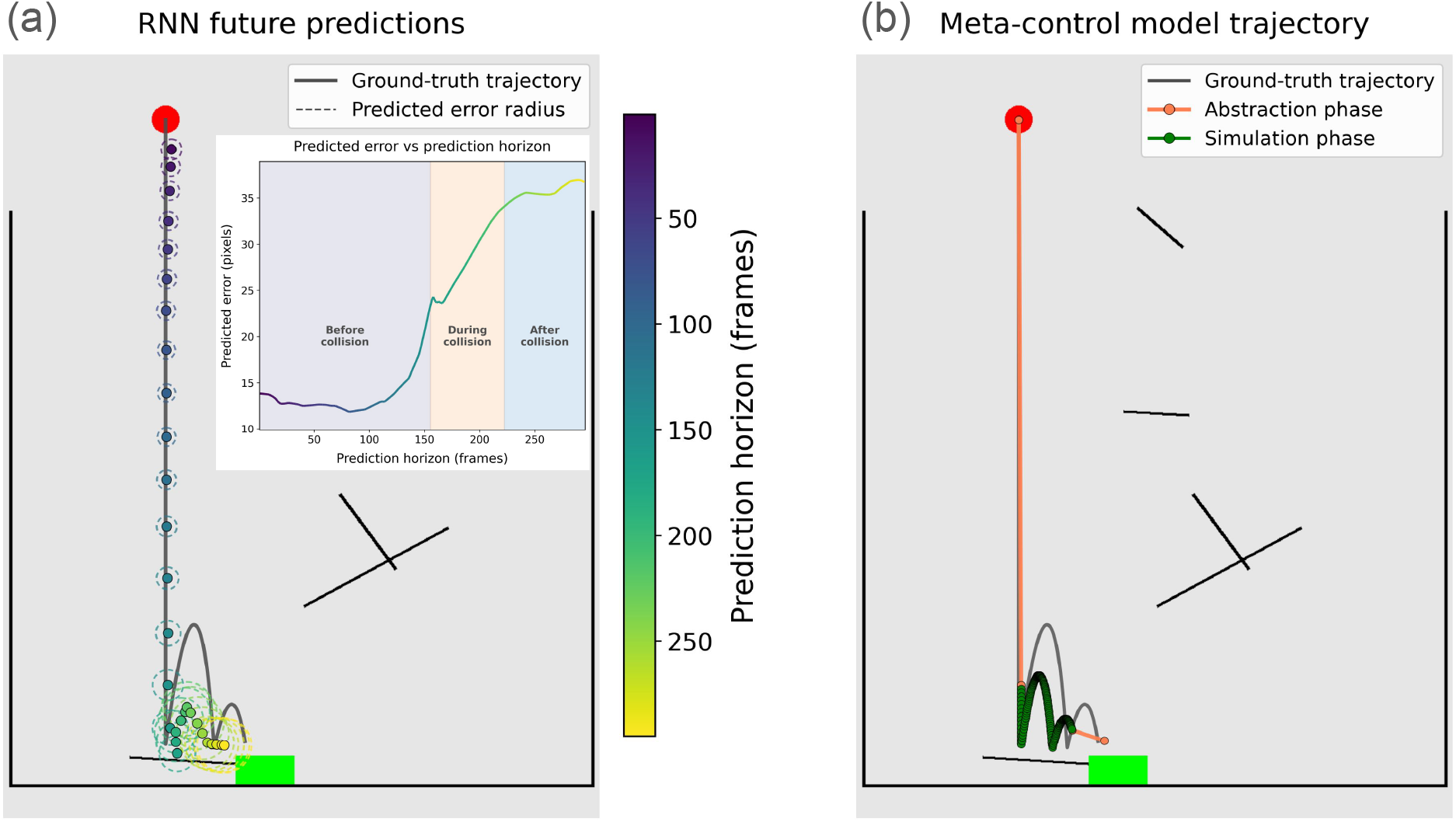
Illustration of RNN and meta-control model predictions. (a) RNN predictions at successive future offsets, with predicted error indicated by dashed circles and the error curve (inset). The color scale indicates the temporal order of predictions across future offsets. The gray line shows the true ball trajectory. (b) Meta-control model prediction of the reasoning trajectory, with orange indicating abstraction phases and green indicating simulation phases.

### The RNN predicts more efficient trajectories in non-straight-path scenes

Given the first 15 frames of each scene, the RNN was queried at successive future offsets to predict the ball’s position, until the predicted ball reached the goal or ground (Fig. 3a). We first examined whether these RNN-based predictions formed more “direct” trajectories than the corresponding ground-truth physical trajectories. Our expectation here was that the RNN would learn shortcuts that can sometimes bypass the ground-truth physical trajectories.

We measured ‘path efficiency’ as the straight-line displacement from the trajectory start to its endpoint, divided by the total distance traveled along the trajectory. Path efficiency ranges from 0 to 1, with a value of 1 indicating a perfectly straight path and lower values indicating increasingly indirect trajectories. We compared the path efficiency of the trajectory created by the RNN-predicted positions across future offsets with that of the ground-truth simulation trajectory, for each scene where the ground truth was a non-straight path. Straight-path scenes were excluded because their ground-truth trajectories have a path efficiency of 1, leaving no possibility for the RNN-predicted trajectory to achieve higher path efficiency.

In non-straight-path scenes, RNN-predicted trajectories had significantly higher path efficiency than the ground-truth trajectories (paired Wilcoxon signed-rank test: median difference = 0.04, *W* = 136.00, *p <* .001). Put more plainly: when the true physical trajectory was indirect (e.g. the ball bounces off of a solid body before continuing to fall downwards), the RNN tended to predict a more direct path toward the endpoint. This is important, as it shows an RNN tasked solely with learning future positions can produce straight-path shortcuts, in line with previously hand-specified such shortcuts used in blended models.

### The RNN predicts its own error in systematic and interpretable ways

We used a linear mixed-effects regression model to examine which factors were associated with the RNN’s estimate of its own prediction error:

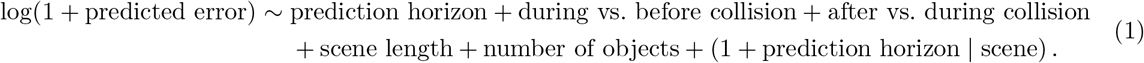

The dependent variable was log(1 + predicted error), where error was measured in pixels. The model included three point-level fixed effects: how far ahead the RNN was predicting, and two contrasts capturing collision stage (during versus before any collision, and after versus during a collision). We additionally included two scene-level fixed effects: total scene length, and the number of objects in the scene. Continuous predictors were standardized. The model allowed each scene to have its own baseline error estimate, and its own relationship between prediction horizon and error.

All five fixed effects were significantly different from zero. The RNN predicted greater error for positions farther into the future, during collisions and especially after collisions, in longer scenes, and in scenes containing more objects (Table 1). These are all reasonable and intuitive factors expected to lead to greater error or uncertainty. Notably, some of these learned error patterns parallel findings in human intuitive physics, where uncertainty increases as physical predictions extend farther and involve additional interactions such as bounces [31].

**Table 1.** Linear mixed-effects model of RNN-predicted error.

| Parameter | $\beta$ | SE | 95% CI | $p$-value |
| --- | --- | --- | --- | --- |
| Prediction horizon | 0.201 | 0.028 | [0.146, 0.256] | < .001 |
| During vs. before collision | 0.209 | 0.005 | [0.198, 0.219] | < .001 |
| After vs. during collision | 0.538 | 0.008 | [0.522, 0.554] | < .001 |
| Scene length | 0.269 | 0.029 | [0.212, 0.325] | < .001 |
| Number of objects | 0.080 | 0.029 | [0.024, 0.136] | .005 |

*Note.* The dependent variable was $\log(1 + \text{predicted error})$ . Continuous predictors were standardized. Collision-stage effects are pairwise contrasts between model-estimated stage means. Marginal $R^2 = .671$ ; conditional $R^2 = .980$ .

The fixed effects alone explained 67.1% of the variance in the RNN’s error estimates, while the full model including random effects explained 98.0% of the variance. These results suggest that the RNN’s predicted error provides a structured estimate of when its position predictions are likely to be less reliable.

### RNN-predicted error is associated with pupil size

The meta-control model uses the RNN-predicted error as a reliability signal for trading off abstraction and simulation. We next asked whether this signal was also related to human performance, and specifically to a pupillary correlate of cognitive effort. Given that pupil dilation is commonly associated with greater cognitive effort [29, 30], we would expect that higher RNN-predicted error be associated with larger mean pupil size. We found that this was indeed the case (*r* = 0.47, *p <* .001; Fig. 4), suggesting that RNN-predicted error may capture aspects of the scene that also influence how much cognitive effort people need to make a judgment.

**Fig 4.**
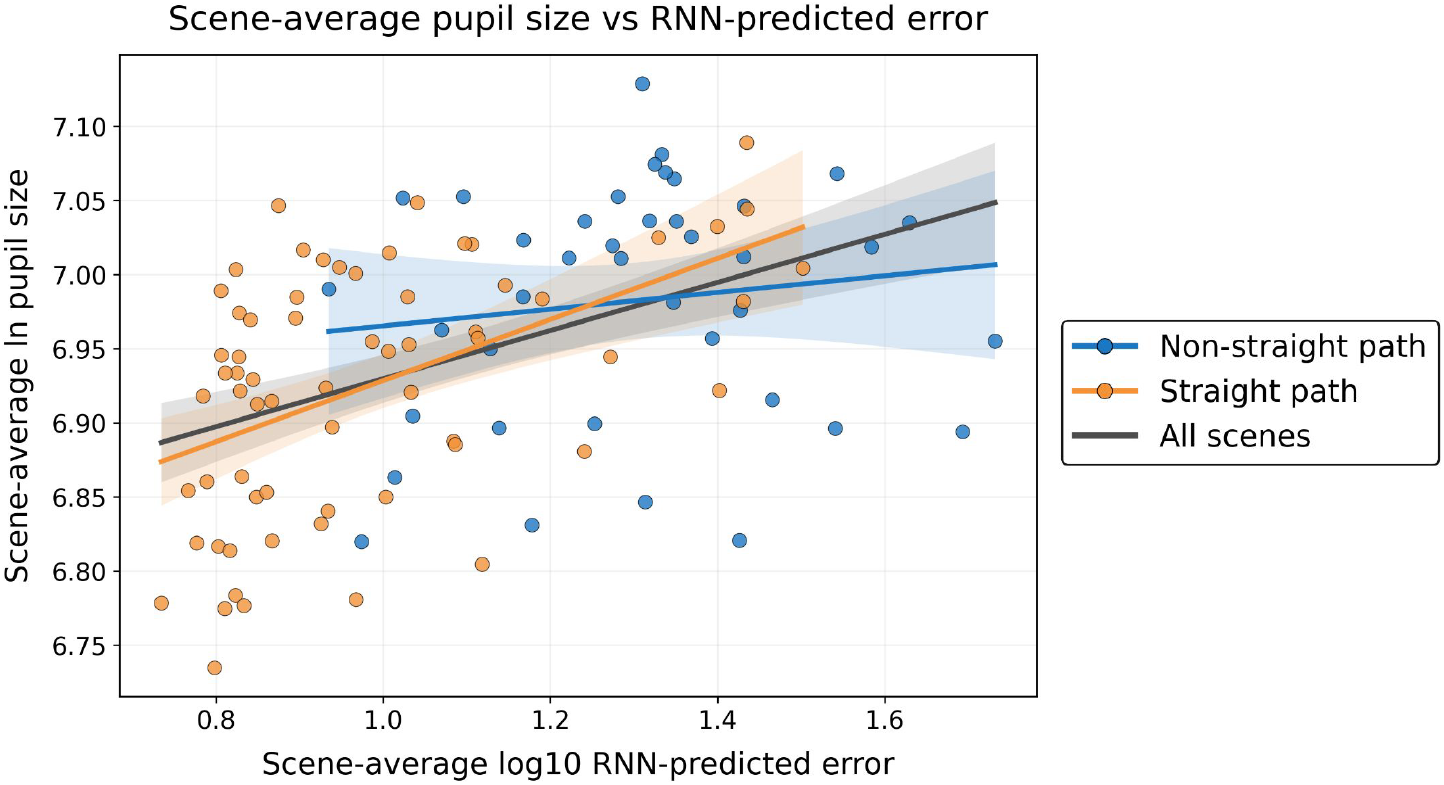
RNN-predicted error is associated with pupil size. Across scenes, higher mean RNN-predicted error was associated with larger mean pupil size during participants’ judgments. When broken down by scene type, this positive relationship was significant for straight-path scenes, but not for non-straight-path scenes. Each point represents a scene; shaded bands indicate 95% confidence intervals for the fitted mean relationships.

It is possible that the relationship we found is rather coarse. That is, the scenes that people saw were either ‘straight path’ (the ball falls directly downwards without hidden obstacles along the way) or ‘non-straight path’ (the ball hits obstacles on the way down and bounces off of them). The correlation between pupil size and predicted error therefore might reflect a binary distinction, ‘difficult’ vs ‘easy’ scenes. To study this in more detail, we examined the relationship between pupil size and RNN-predicted error separately for straight-path and non-straight-path scenes. We found that the relationship was significant for straight-path scenes (*r* = 0.48, *p <* .001), but not for non-straight-path scenes (*r* = 0.14, *p* = 0.41; Fig. 4). One possible interpretation of this result is that mean pupil size during judgment primarily reflects whether simulation is engaged for a given scene. At the scene level, higher mean RNN-predicted error may increase the likelihood of engaging simulation in straight-path scenes. In non-straight-path scenes, however, simulation may already be engaged with near-ceiling probability, so further increases in mean predicted error would have little additional effect on pupil size.

### The meta-control model captures human behavior and eye-movement patterns

Having shown that the RNN provides a direct predictive abstraction and a structured estimate of its own error, we next examined whether the meta-control model, which uses this error signal to regulate abstraction and simulation, captures key patterns in people’s response time, accuracy, and eye movements.

#### Response-time patterns

The meta-control model captured the human response-time pattern well (Fig. 5). For scenes with non-straight trajectories, longer ground-truth simulation times were associated with longer response times in humans and longer model run times (OLS slopes: human, *β* = 0.12, *t*(44) = 2.98, *p* = .005; model, *β* = 0.87, *t*(44) = 6.24, *p <* .001). In contrast, these relationships were significantly weaker for straight-path scenes (OLS interactions: human, *β* = −0.09, *t*(44) = −1.79, *p* = .040; model, *β* = −0.50, *t*(44) = −2.74, *p* = .004). Thus, both people and the model showed a stronger relationship between reasoning time and ground-truth simulation time for non-straight trajectories, where a simple shortcut is less likely to suffice and simulation is more likely to be engaged, and a weaker relationship when the ball could follow a straight path to the goal.

**Fig 5.**
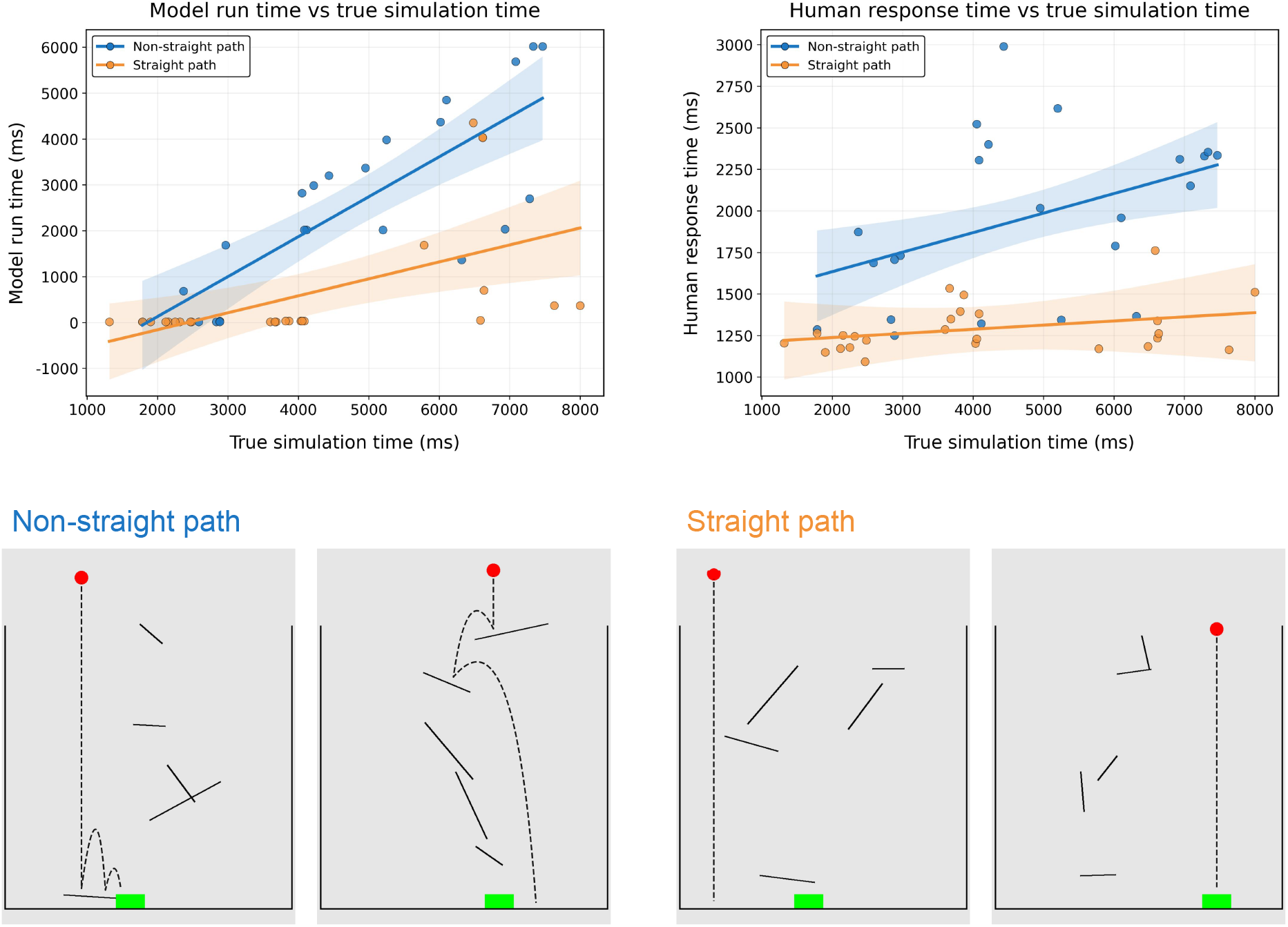
The meta-control model captures the human response-time pattern. Top row: Human response times and model run times showed the same pattern: both showed strong positive associations with ground-truth simulation time for non-straight trajectories, but this relationship was weaker when the ball could follow a straight path to the goal. Each point represents a scene; shaded bands indicate 95% confidence intervals for the fitted mean relationships. Bottom row: Example scenes from the straight-path and non-straight-path conditions. The dashed black line indicates the ball’s ground-truth trajectory under pure simulation.

#### Accuracy patterns

Human accuracy followed a non-monotonic pattern as a function of ground-truth simulation time (Fig. 6), replicating the key finding of [12]. These results were qualitatively captured by the meta-control model (Fig. 6). Consistent with this qualitative match, human accuracy was significantly higher in scenes where the meta-control model predicted correctly than in scenes where it predicted incorrectly (Mann–Whitney *U* test: median difference = 0.50, *U* = 20.00, *p <* .001).

**Fig 6.**
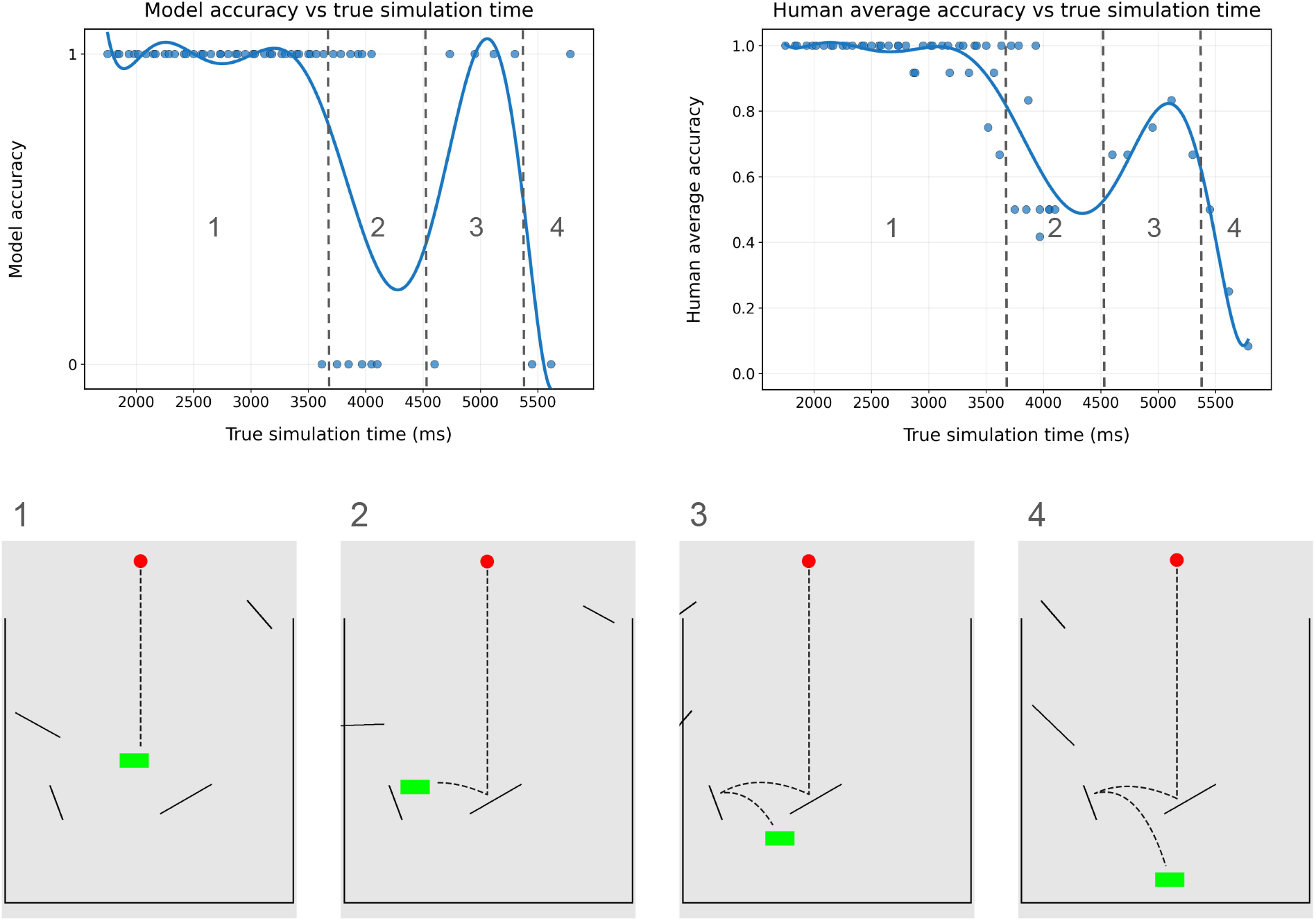
The meta-control model captures the human accuracy pattern well. Top row: The meta-control model captured the nonlinear relationship between human accuracy and ground-truth simulation time. For visualization, the relationship was fit with an eighth-order polynomial curve. Participant accuracy starts off near ceiling for the first group of scenes in Section 1, in which the ball follows a short, straight path. Accuracy then takes a sharp dip for the second group of scenes in Section 2, before rising slightly in Section 3, and dipping again in Section 4. These results are qualitatively captured by the meta-control model. Bottom row: Example scenes from each of the Sections 1–4. The dashed black line indicates the ball’s ground-truth trajectory under pure simulation.

#### Eye-movement signatures of abstraction and simulation

We next asked whether the abstraction and simulation phases predicted by the meta-control model corresponded to distinct patterns of human eye movements. Using the NSLR-HMM method [32], we classified gaze samples into saccades, smooth pursuits, fixations, and post-saccadic oscillations. We focused on saccades and smooth pursuits because they have natural interpretations as signatures of abstraction and simulation, respectively. Saccades are consistent with rapid visuospatial abstraction, in which gaze shifts or jumps toward a predicted location, whereas smooth pursuit can reflect continuous tracking of perceived or imagined motion and is therefore naturally associated with step-by-step simulation.

We used the meta-control model to identify abstraction and simulation phases along the reasoning trajectory (Fig. 3b). We then calculated the proportions of nearby human gaze samples classified as saccades or smooth pursuits within each model-predicted phase. To identify gaze samples associated with each trajectory segment, we tested perpendicular-distance thresholds from 50 to 120 pixels in 10-pixel increments. The results were qualitatively consistent across thresholds (Fig. 7a,b). At the 120-pixel threshold reported here, participants showed a significantly higher proportion of saccadic eye movements during model-predicted abstraction phases than during simulation phases (within-subject bootstrap test: median difference = 0.08, 95% CI [0.02, 0.12], *p* = .004; Fig. 7a). Conversely, smooth-pursuit movements were significantly more common during model-predicted simulation phases than during abstraction phases (within-subject bootstrap test: median difference = 0.02, 95% CI [0.00, 0.04], *p* = .012; Fig. 7b). These results provide evidence that the two reasoning modes identified by the model correspond to distinct patterns of human eye movements.

**Fig 7.**
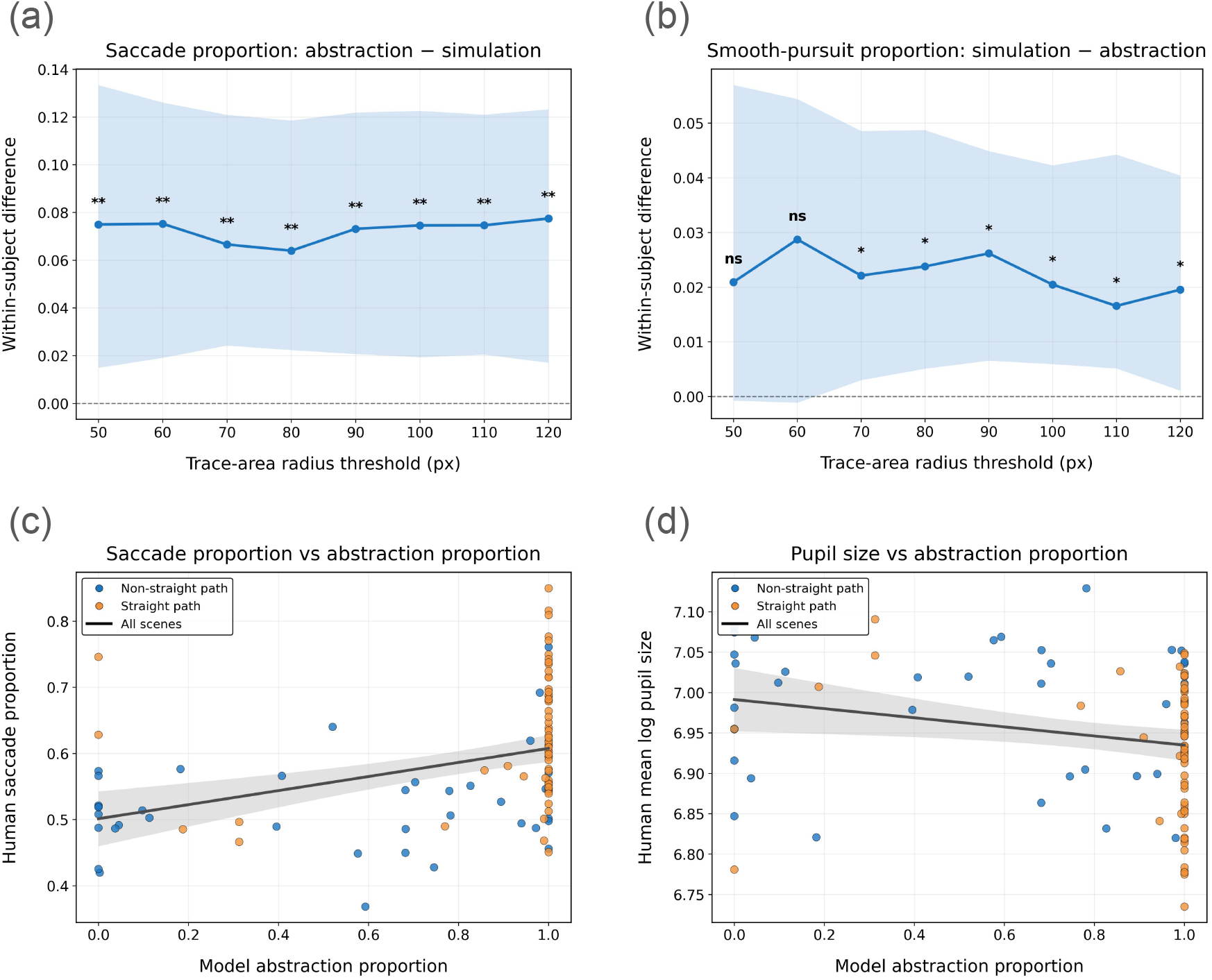
The meta-control model captures the human eye-movement patterns well. (a)-(b) Participants showed a higher proportion of saccadic eye movements during model-predicted abstraction phases than during simulation phases, and a higher proportion of smooth-pursuit eye movements during simulation phases than during abstraction phases. This pattern was consistent across a range of perpendicular-distance thresholds used to associate gaze samples with each trajectory segment. Each point represents the median within-subject difference, and the shaded area indicates the 95% confidence interval from the bootstrap. (c)-(d) At the scene level, higher model-predicted abstraction proportion was associated with a higher average proportion of saccades and smaller average pupil size in the human eye-gaze data. Abstraction proportion was defined as the total length of abstraction segments divided by the total reasoning-trajectory length within each scene. Each point represents a scene; shaded bands indicate 95% confidence intervals for the fitted mean relationships.

At the scene level, model-predicted abstraction proportion was computed as the total length of abstraction segments divided by the total length of the reasoning trajectory within each scene. A higher model-predicted abstraction proportion was associated with a higher average proportion of saccades in human eye-gaze data (*r* = 0.40, *p <* .001; Fig. 7c) and smaller average pupil size (*r* = −0.23, *p* = .016; Fig. 7d). These associations are consistent with model-predicted abstraction phases involving less continuous visual tracking and lower cognitive effort than simulation.

### The meta-control model improves correspondence with human response time and eye movements relative to the earlier blended model

As mentioned in the Introduction, our meta-control model improves conceptually on the ‘blended model’ used in our previous work [12]. This blended model combines a partial simulation that simulates the scene forward by a small number of steps with a simple abstraction that projects the ball forward along its current direction. At each moment, the model computes and compares the predictions of partial simulation and linear projection: when they are sufficiently similar, the model adopts the abstraction update, whereas when they diverge, it relies on simulation. By contrast, our meta-control model requires no partial simulation and learns abstractions rather than having them hand-coded. But, is the meta-control model also in better correspondence with human data?

### Stronger correspondence with human response time

The relationship between model run time and human response time was significantly stronger for the meta-control model than for the blended model (Williams test: Δ*r* = 0.19, *t*(45) = 2.54, *p* = .014; Fig. 8a). Time measures were log-transformed to reduce heteroskedasticity.

**Fig 8.**
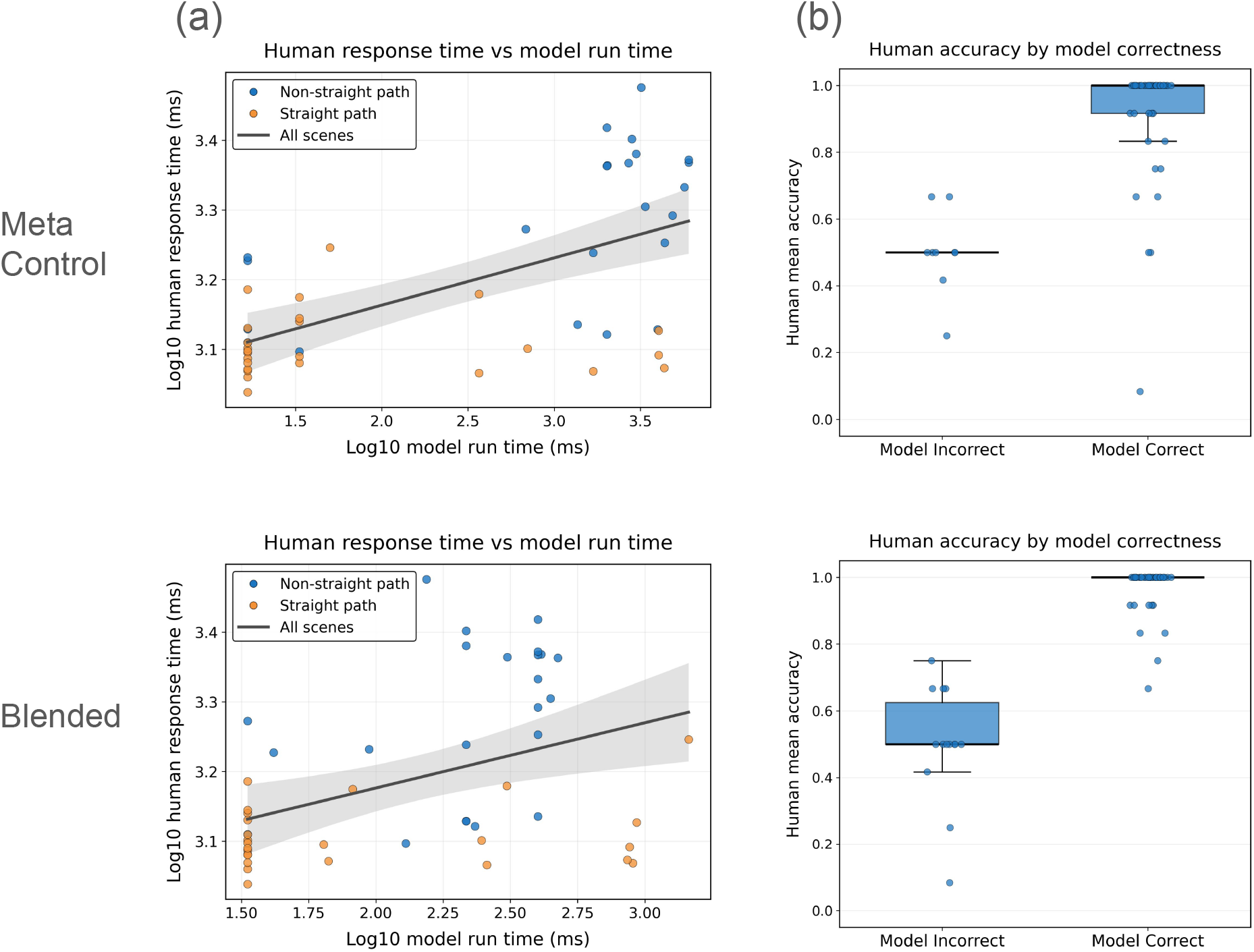
The meta-control model better captures human response time than the blended model. (a) The relationship between model run time and human response time was stronger for the meta-control model than for the earlier blended model. Each point represents a scene; shaded bands indicate 95% confidence intervals for the fitted mean relationships. Time measures were log-transformed to reduce heteroskedasticity. (b) In both models, human accuracy was higher in scenes that the model predicted correctly than in scenes that it predicted incorrectly.

#### Both models capture human accuracy patterns

The meta-control and blended models showed similar qualitative correspondence with human accuracy. Both captured the nonlinear relationship between human accuracy and ground-truth simulation time. In addition, for both models, human accuracy was higher in scenes that the model predicted correctly than in scenes that it predicted incorrectly (Mann–Whitney *U* test: meta-control, median difference = 0.50, *U* = 20.00, *p <* .001; blended, median difference = 0.50, *U* = 3.00, *p <* .001; Fig. 8b).

#### Stronger separation of eye-movement signatures

We next asked whether the meta-control model’s abstraction-versus-simulation classifications better predicted the corresponding human eye-movement signatures than those of the blended model. We included gaze samples that were either a saccade or a smooth pursuit and that were uniquely classified as abstraction or simulation by both models at each distance threshold. For each model, we fit a logistic regression for gaze samples *i* within participants *j*:

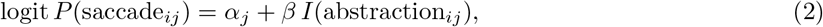

where *α*_*j*_ is a participant-specific intercept capturing individual differences in baseline saccade probability.

We evaluated predictive performance using 10-fold cross-validation at the scene level. Scenes were partitioned into 10 folds; in each iteration, models were fit by maximum likelihood on gaze samples from nine folds and tested on the remaining fold. Performance was quantified by held-out log-likelihood, with higher values indicating better prediction of whether each gaze sample was a saccade or smooth pursuit.

Across distance thresholds, the meta-control model’s phase classifications generally predicted human saccade-versus-smooth-pursuit behavior better than those of the blended model (Fig. 9). At the 120-pixel threshold reported here, the meta-control model showed a small but reliable predictive advantage (scene-level sign-flip test: Δ*LL* = 0.002, *p* = .004; bootstrap: 95% CI [0.001, 0.004]). This advantage indicates that the distinction between abstraction and simulation in the meta-control model more closely corresponded to the separation of saccadic and smooth-pursuit eye movements observed in people.

**Fig 9.**
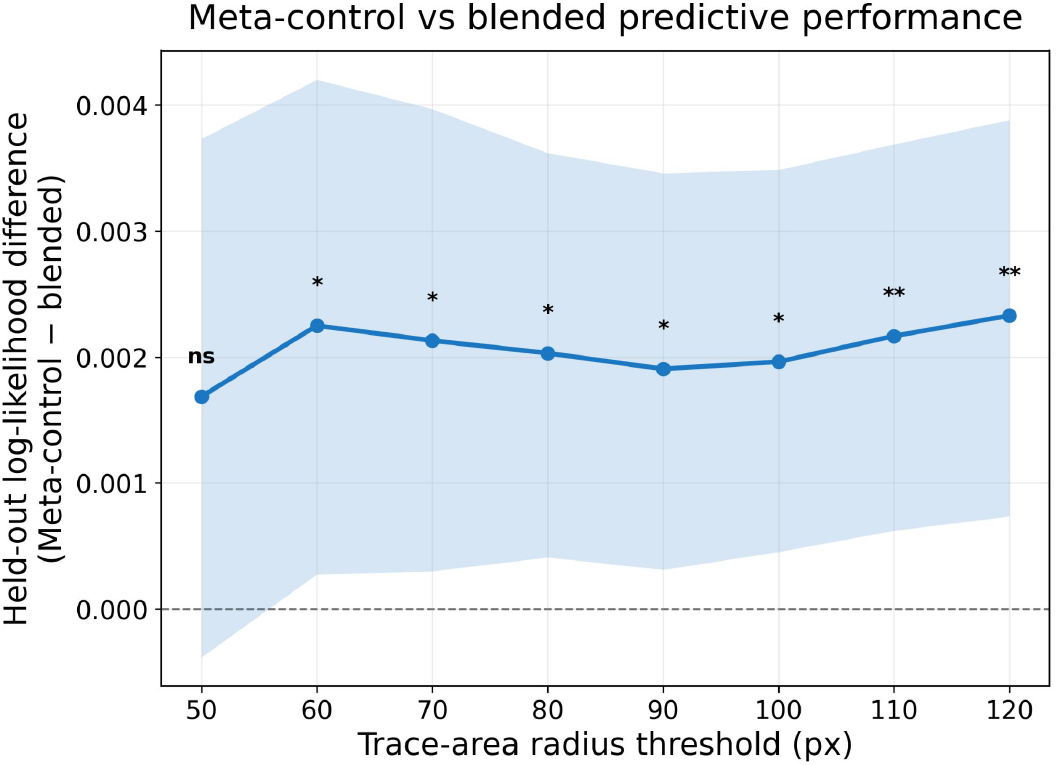
The meta-control model better captured human eye-movement patterns than the blended model. The meta-control model’s abstraction-versus-simulation classifications predicted human saccade-versus-smooth-pursuit behavior better than those of the blended model. This advantage was consistent across a range of perpendicular-distance thresholds used to associate gaze samples with trajectory segments. Each point shows the mean difference in held-out log-likelihood between the two models; shaded bands show 95% confidence intervals from a scene-level bootstrap.

## Discussion

Our results provide converging evidence that intuitive physical reasoning can be captured by a reliability-guided coordination of abstraction and simulation. We implemented a physical-prediction RNN that learned a direct abstraction, as well as an estimate of when its predictions were likely to be less reliable. We then constructed a meta-control model that uses the error estimate to trade off abstraction and simulation. We found that this meta-control model captured human response-time and accuracy patterns, while its predicted phases corresponded to distinct eye-movement signatures. The meta-control model improves conceptually on an earlier blended model [12], and also showed stronger correspondence with human response times and eye movements.

Our findings suggest that the alternative to detailed simulation need not take the form of a predefined heuristic. Instead, predictive experience can produce an abstraction that bypasses intermediate physical states, together with an estimate of when that abstraction can be trusted. The stronger human-model correspondence relative to the blended model suggests that learned predictive reliability may provide a useful signal for determining when detailed simulation is needed.

Several limitations leave important questions for future work. First, eye movements and pupil size provide converging process-level evidence, but neither uniquely identifies the underlying computation. Smooth pursuit can reflect multiple visual and cognitive processes, including motion tracking, attention, prediction, and memory, and therefore does not uniquely identify simulation [33, 34]. Pupil size is likewise influenced by several processes beyond cognitive effort [35]. More generally, similar patterns can come about from different cognitive mechanisms [36]. Future work could experimentally encourage participants to use simulation or abstraction, and test whether the two reasoning modes produce the predicted eye-movement and pupil signatures. This would provide a more direct test of whether these oculomotor patterns reflect distinct reasoning processes.

Second, although the abstraction mechanism and its error estimate are learned, the meta-control policy is still specified by the modeler. A fixed threshold determines when the model switches to simulation, and the simulator itself is provided in advance. Previous work shows that people can learn which strategies or computations to select through experience [37], and recent models have formalized dynamic allocation of computation according to its expected usefulness [38–41]. A natural extension would therefore be to train the model on human behavioral data to learn both the reasoning strategies and their allocation, and test whether a reliability-guided combination of abstraction and simulation emerges without being specified in advance.

Third, the present study considered only one prediction task within a restricted family of physical scenes. People can reason about the dynamics of various everyday scenes across substantially different scenarios and inference demands, suggesting that an important test is whether the learned abstraction and meta-control policy generalize in the same way [42, 43]. Future work could test the model across different physical domains, such as stability, containment, or multi-object interactions, and ask whether an abstraction learned for one type of physical prediction transfers to another or must be adapted to the new task.

Finally, our measures invite future research on the neural mechanisms underlying abstraction, simulation, or switching between them. Recent work has identified invariant representations of physical properties and predicted future states in a frontoparietal “physics network” [44–46], while the dorsal anterior cingulate cortex has been proposed to regulate when and how intuitive physics processes are engaged [47]. Combining the present paradigm with neural measures such as fMRI could test whether model-predicted abstraction and simulation phases correspond to distinct neural dynamics, and whether predicted error or switching points are represented in regions that support or regulate physical inference.

Our results advance the understanding of intuitive physics by showing how people balance computational efficiency and accuracy through reliability-guided switching between abstraction and simulation. More broadly, they suggest a simple answer to how people can make useful physical predictions without incurring the cognitive demands of simulating every moment in detail: learning when it is safe to rely on abstractions. Visual experience can give rise to predictive abstractions that jump over intermediate physical states together with estimates of their reliability. Rather than relying on a single fixed strategy, people may reason efficiently about physical events by flexibly deciding when to trust an abstraction and when to engage mental simulation.

## Materials and Methods

The behavioral and eye-tracking data analyzed here were collected in an earlier study [48].

### Participants

Participants (*N* =18) were recruited via Harvard University’s Study Pool. The participants were undergraduate students, graduate students, or post-doctoral researchers. All participants provided informed consent, and the study was approved by the Harvard University Committee on the Use of Human Subjects (IRB22-0349). Participants were compensated for their time ($15.00/hr). Six participants were excluded from the analysis due to early issues with data collection and data corruption, leaving *N* =12 participants for analysis.

### Procedure

Participants viewed physical scenes containing a ball, a goal, and a variable number of obstacles termed slides (Fig. 10). Responses were made using a three-button controller with “Yes,” “No,” and “Next” buttons; “Yes” and “No” assignments were randomized across participants, while “Next” was always the middle button. Participants completed the task in a silent room with their heads positioned on a chin rest 55 cm from the screen. The right eye was tracked using an SR Research EyeLink 1000 Plus at 1000 Hz and calibrated using a 13-point procedure before the experiment.

**Fig 10.**
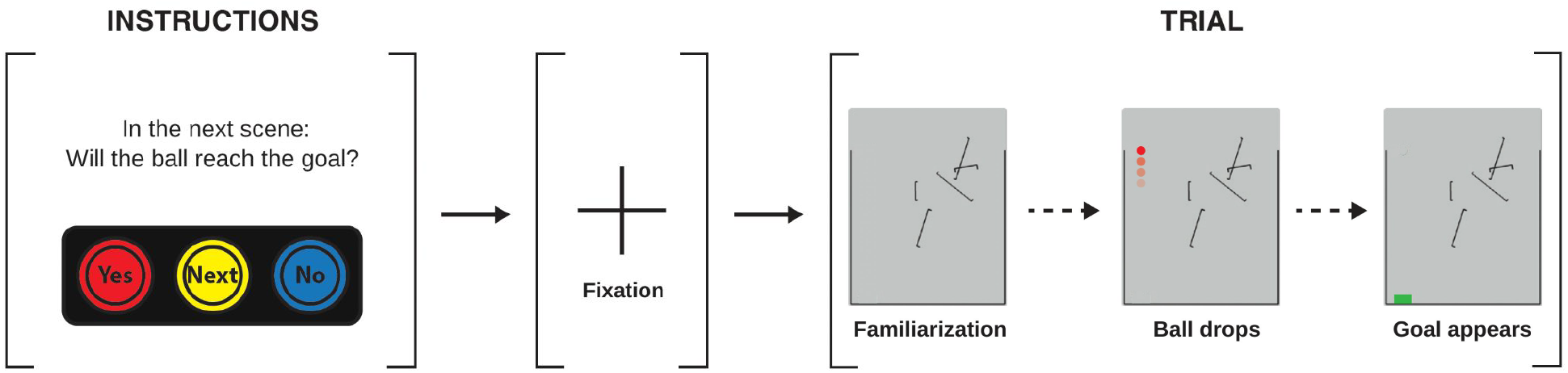
Experiment procedure. Participants sit in front a screen that displays a falling ball, and use different colored button to indicate their response. Participants are first reminded of task instructions and controller button mappings. Participant eye movements are recorded throughout the task using an eye tracker. Participants begin with a short fixation period, before proceeding to experimental trials. Each trial consists of three phases. First, participants familiarize themselves with a given scene. They are shown black lines that act as impenetrable obstacles that a ball cannot pass through. Second, a ball (red circle) appears and begins to fall, gradually fading out over 1 second. Participants are instructed to imagining the ball continuing to fall after it disappears. Third, a goal (green rectangle) appears at a random time, and participants respond as quickly as possible to indicate whether the ball would have hit the goal.

#### Familiarization

Participants completed three familiarization blocks of increasing difficulty. The first familiarized participants with the controller by asking them to report whether a visible ball trajectory ended in a collision with the goal. In the second, the ball fell for 1 second (60 simulation steps) and then disappeared, after which participants judged whether it would ultimately collide with the goal. The final familiarization block mirrored the main experiment: participants first inspected the slides at their own pace, then viewed the ball falling for 1 second before disappearing, and finally judged as quickly as possible whether the ball would have collided with the subsequently presented goal.

#### Experiment

Stimuli were presented using a pseudo-Latin square design with four treatments and were randomized within treatment across participants. On each block, participants first viewed the slides alone and inspected the scene at their own pace. After pressing “Next,” the ball appeared, fell for 1 second, and disappeared. The goal was then presented, and participants responded as quickly as possible whether the ball would have collided with it. This block structure was designed to separate eye-gaze behavior associated with visual scanning from behavior associated with reasoning about the ball’s trajectory.

### Stimuli

In total, our eye-tracking experiment contained 106 stimuli, all of which were drawn directly from [12], with only the modifications needed to align them with the block structure of the eye-tracking experiment. Each stimulus showed a single scenario containing a ball (a red circle), a goal (a green rectangle), and slides (narrow, rigid black rectangles) on a white background. All stimuli were created using the 2D rigid-body physics engine Pymunk (https://www.pymunk.org) and rendered as videos at 60 frames per second. A random subset was flipped about the horizontal axis before presentation.

The stimuli were designed to distinguish pure simulation, pure abstraction, and blended models based on their predicted response times and accuracy across scenes. The 48 stimuli in Experiment 1 were designed to distinguish the models based on their response-time predictions, whereas the 58 stimuli in Experiment 2 were designed to distinguish the models based on their accuracy predictions. All 106 stimuli from both experiments were included in the eye-movement analyses.

### Modeling

#### RNN model for future-position and error prediction

##### Architecture

We developed a recurrent neural network (RNN) inspired by the Index-and-Track (InT) circuit [49], adapting its local recurrent processing to predict future positions and estimate prediction error. The model receives 15 consecutive RGB frames, each resized to 100 × 128 pixels. Each frame is processed by three convolutional layers: a 5 × 5 convolution with 32 channels and stride 2, a 3 × 3 convolution with 64 channels and stride 2, and a 3 × 3 convolution with 96 channels, with ReLU activations after each layer. The recurrent component maintains a 96-channel spatial state and applies a depthwise 3 × 3 recurrent convolution, with one recurrent kernel per channel. At each time step, a candidate state is computed from the current visual features and the locally transformed previous state, and a learned 1 × 1 convolution produces an update gate that determines how much of the candidate state replaces the previous state. After all 15 frames are processed, learned spatial attention pools the final recurrent state into a 96-dimensional representation, which is transformed by a two-layer multilayer perceptron (MLP) into a 192-dimensional scene representation.

To make a prediction, the model additionally receives the requested future offset. The normalized offset *τ* is encoded using *τ, τ* ^2^, sin(*πτ* ), and cos(*πτ* ) to provide a richer nonlinear basis; a two-layer MLP then transforms these features into a 32-dimensional time representation. The scene and time representations are concatenated and passed through a shared decoder with hidden dimensions 192 and 96. Two output heads then predict the ball’s two-dimensional position and the magnitude of the corresponding position-prediction error. The error output is constrained to be positive using a softplus transformation. Given an encoded visual history, each future position and corresponding error estimate is predicted independently at the requested offset, rather than autoregressively from earlier predictions. Thus, predicting a distant future state does not require first predicting every intermediate state. The complete model contains 214,820 trainable parameters.

##### Training

The training set comprised 6,000 scenes generated by randomizing the number and positions of the objects present in the 106 scenes used in the human experiment. After the ball reached the goal or ground, its final position was held fixed for 1 second (60 frames) to provide post-event prediction targets. RGB frames were resized to 100 × 128 pixels and scaled to the interval [0, 1]. Ball positions were represented in the original 800 × 1024 pixel coordinate system, with horizontal and vertical coordinates normalized by image width and height, respectively.

For each training scene, 20 windows of 15 consecutive frames were randomly sampled per epoch, with all input frames occurring before the terminal frozen period. Across 6,000 training scenes, this yielded 120,000 training windows per epoch. For an input window ending at frame *t*, the model predicted the ball’s position and corresponding prediction-error magnitude at every valid future offset *k*, beginning at *k* = 0 at the final observed frame and continuing through the end of the recorded scene, including the post-event frozen period. Future offsets were normalized by the maximum prediction horizon across the training and experimental scenes.

For training window *s*, let *t*_*s*_ denote the final observed frame and *K*_*s*_ the set of valid future offsets. For each *k* ∈ *K*_*s*_, let 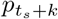 denote the ground-truth normalized ball position and 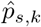 the predicted position. The position loss was defined as

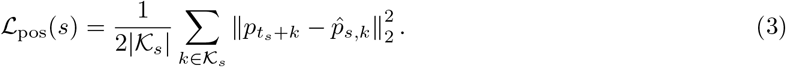

The target for the error head was the Euclidean magnitude of the normalized position-prediction error,

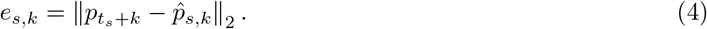

When computing the error-prediction loss, *e*_*s,k*_ was treated as a detached target, so gradients from this loss did not propagate through the position prediction used to compute *e*_*s,k*_.

The error-prediction loss was

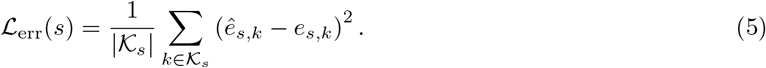

The total training loss was

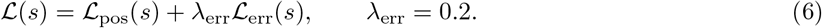

Because scenes differed in the number of valid future offsets, target sequences were padded and masked so that only valid offsets contributed to the loss. Losses were first averaged over valid offsets within each training window and then averaged equally across windows within each minibatch.

The model was trained for 150 epochs using AdamW with a batch size of 32, a learning rate of 10^−4^, and weight decay of 10^−5^. Gradient norms were clipped at 1.0. No human behavioral or eye-tracking measures were used to train the RNN.

##### Inference

At test time, the model received the first 15 frames of each experimental scene and was queried at successive future offsets to generate position and prediction-error estimates until the predicted ball reached the goal or ground. Normalized position predictions were converted back to the original 800 × 1024 pixel coordinate system by scaling the horizontal and vertical coordinates by 800 and 1024, respectively. Predicted error magnitude was converted to an approximate pixel scale by multiplying by the mean of these coordinate scales, 912.

#### Meta-control model

##### Algorithm

The meta-control model uses the RNN as a fast abstraction mechanism and Pymunk as a physics simulator (Algorithm 1). Starting from the initial 15-frame RNN history, the controller queries increasingly distant abstraction endpoints at offsets *A*, 2*A*, 3*A*, … and accepts each abstraction as long as its predicted endpoint error remains below the threshold *τ* . If the predicted error exceeds *τ*, the controller switches to Pymunk to generate increasingly long simulation segments of length *S*, 2*S*, 3*S*, … until the final 15 simulated frames provide a new RNN history for which the predicted error of the next *A*-frame abstraction falls below *τ*, at which point the controller updates the RNN history and resumes abstraction.

##### Parameters

The meta-control model has three parameters: the fixed abstraction span *A*, the simulation-length increment *S*, and the predicted-error threshold *τ* . We performed a grid search over these parameters and selected the combination that produced the strongest correlation between model runtime and human response time across scenes. Both measures were log-transformed to reduce heteroscedasticity before computing the correlation. The parameters used for all subsequent meta-control model evaluations were *A* = 150, *S* = 20, *τ* = 22.

#### Blended model

For comparison, we used the same blended model as in our earlier work [12]. In this model, physical reasoning combines two processes: a partial simulation that simulates the scene forward by a small number of steps and a simple abstraction that projects the ball forward along its current direction. At each moment, the model computes a possible next state using *N* steps of partial simulation and, in parallel, a linear projection of length *D*. It then compares the predictions of the two processes: when their cosine similarity is at least *E*, the model adopts the abstraction update; otherwise, it relies on simulation. This real-time trade-off allows the model to short-circuit simulation and conserve computational resources when *D* is greater than *N* . We retained the original parameter values: *N* = 5, *D* = 75, *E* = 0.9.

### Analyses

#### RNN trajectory efficiency

To examine the structure of the RNN’s predicted trajectories, we computed path efficiency as the straight-line displacement from the trajectory start to its endpoint divided by the total distance traveled. RNN-predicted and ground-truth path efficiencies were compared across non-straight-path scenes using a paired, two-sided Wilcoxon signed-rank test. Straight-path scenes were excluded because their ground-truth path efficiency was 1.

#### Factors associated with RNN-predicted error

To examine factors associated with the RNN’s predicted error, we fit a linear mixed-effects model with log(1 + predicted error) as the dependent variable. Fixed effects included standardized prediction horizon, two planned contrasts comparing during versus before the first collision and after versus during the first collision, standardized scene length, and standardized number of objects. Later collision periods were grouped into the post-collision stage. The model included a random intercept and random slope for prediction horizon by scene and was fit by maximum likelihood. Fixed-effect significance was assessed using Wald tests, and marginal and conditional *R*^2^ quantified variance explained by the fixed effects and by the full model, respectively.

##### Algorithm 1

Meta-Control Algorithm

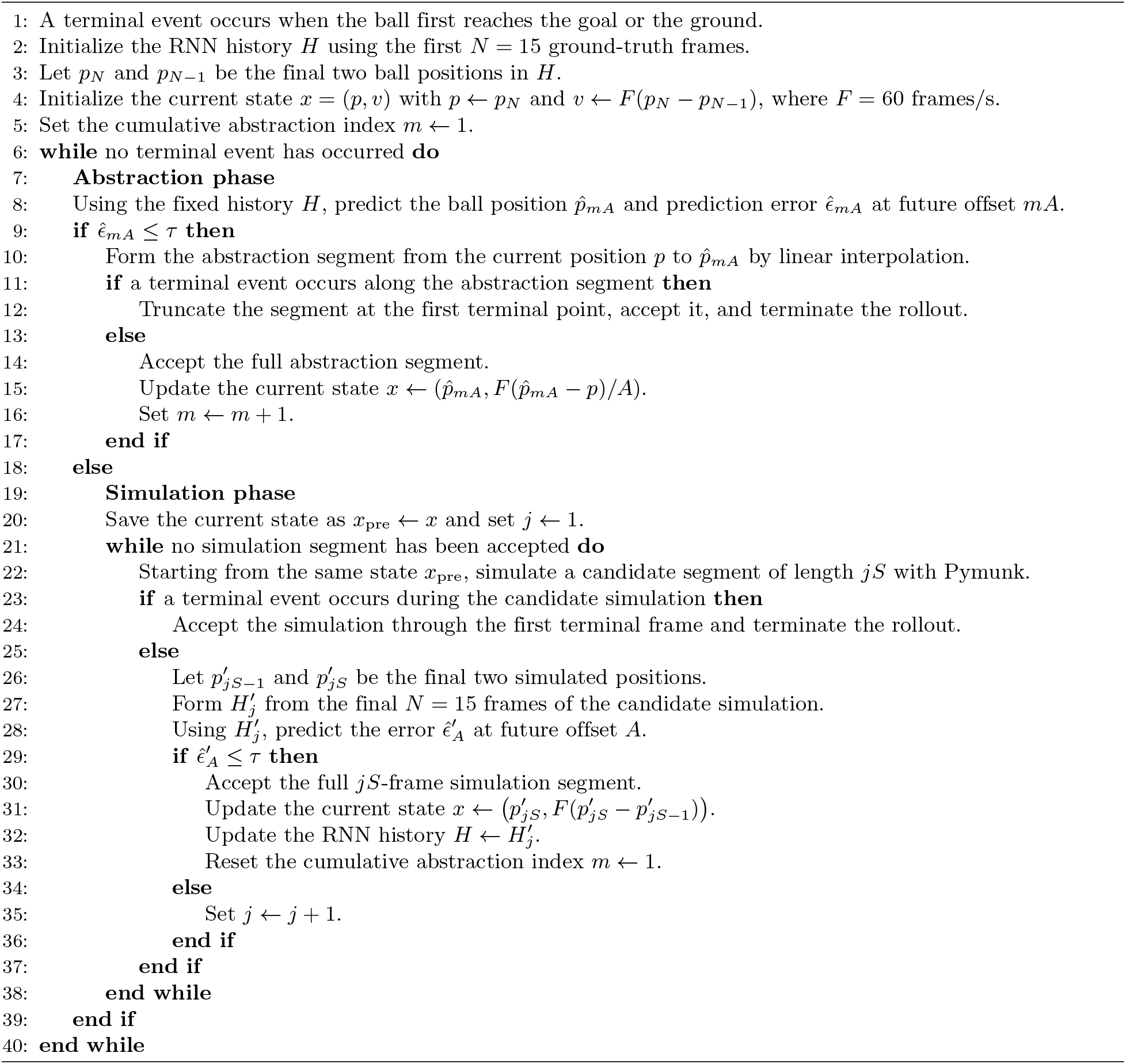

#### RNN-predicted error and pupil size

To relate RNN-predicted error to human pupil size, pupil size was log transformed and averaged first within each participant and scene and then across participants. RNN-predicted error was averaged across prediction horizons within each scene and log_10_ transformed. We calculated Pearson correlations between scene-average RNN-predicted error and pupil size across all scenes and separately for straight-path and non-straight-path scenes. Ordinary least squares (OLS) fits with 95% confidence intervals were used for visualization.

#### Response-time analyses

To compare reasoning-time patterns in humans and the meta-control model, we analyzed Experiment 1 scenes using separate OLS regressions for human response time and model run time. Each model included centered ground-truth simulation time, path type, and their interaction, with non-straight-path scenes as the reference group. Group-specific slopes were tested against zero using two-sided tests. Because our prediction was that the relationship with simulation time would be weaker for straight-path scenes, the interaction was tested one-sided. To compare the meta-control and blended models directly, human response time and model run time were log-transformed to reduce heteroskedasticity, and their Pearson correlations were compared on the same set of Experiment 1 scenes using a two-sided Williams test for dependent overlapping correlations.

#### Accuracy analyses

Human accuracy in model-correct and model-incorrect Experiment 2 scenes was compared using two-sided Mann–Whitney *U* tests. The eighth-order polynomial curves relating accuracy to ground-truth simulation time were used only for visualization.

#### Eye-movement signatures of abstraction and simulation

For the eye-movement analyses, we associated each gaze sample with model-predicted abstraction and simulation segments using its shortest distance to each trajectory segment. We repeated the analysis using distance thresholds from 50 to 120 pixels in 10-pixel increments, with 120 pixels as the primary threshold. For each participant and threshold, we calculated the proportion of saccades and smooth pursuits among gaze samples associated with abstraction and simulation. Samples falling within threshold of both phase types were included in both phase pools.

We then calculated within-participant differences in saccade proportion between abstraction and simulation and in smooth-pursuit proportion between simulation and abstraction. Inference was based on 5,000 bootstrap resamples of participants with replacement. For each resample, we calculated the median within-participant difference; 95% confidence intervals were defined by the 2.5th and 97.5th percentiles, and two-sided bootstrap *p*-values were calculated as twice the smaller empirical tail probability relative to zero.

#### Scene-level abstraction and eye-movement measures

At the scene level, model-predicted abstraction proportion was defined as the summed length of abstraction segments divided by the summed length of all abstraction and simulation segments. Human saccade proportion and log-transformed pupil size were first averaged within participant and scene and then across participants. Pearson correlations were used to relate scene-level abstraction proportion to each human eye measure.

#### Comparison of eye-movement predictions between models

To compare how well the meta-control and blended models separated the two eye-movement signatures, we retained gaze samples that were either a saccade or a smooth pursuit and received a unique abstraction or simulation classification from both models. For each model, we fit a Bernoulli logistic regression predicting saccade versus smooth pursuit from the model-predicted reasoning phase, with participant-specific intercepts included as fixed effects.

Predictive performance was evaluated using 10-fold cross-validation grouped by scene, such that all gaze samples from a held-out scene were excluded from training and both models were evaluated on identical folds and observations. Performance was measured using held-out Bernoulli log-likelihood. We first averaged the meta-control-minus-blended log-likelihood difference within each scene and then averaged across scenes. A 95% confidence interval was obtained from 20,000 bootstrap resamples of scenes with replacement. Because the two models were evaluated on the same scenes, significance was assessed separately using a paired scene-level sign-flip randomization test under the null hypothesis of no systematic difference between models. The sign-flip test used 20,000 random sign assignments to the scene-level differences to estimate the *p*-value. The same analyses were repeated across the full range of distance thresholds as a robustness check.

## Acknowledgments

This work was supported by the Kempner Institute for the Study of Natural and Artificial Intelligence, a Polymath Award from Schmidt Sciences, and the Department of Defense MURI program under Army Research Office grant W911NF-23-10277.

## References

1. Battaglia PW, Hamrick JB, Tenenbaum JB. Simulation as an engine of physical scene understanding. Proc Natl Acad Sci U S A. 2013 Nov;110(45):18327–32. doi:10.1073/pnas.1306572110.

2. Fischer J, Mikhael JG, Tenenbaum JB, Kanwisher N. Functional neuroanatomy of intuitive physical inference. Proc Natl Acad Sci U S A. 2016 Aug;113(34):E5072–81. doi:10.1073/pnas.1610344113.

3. Ahuja A, Sheinberg DL. Behavioral and oculomotor evidence for visual simulation of object movement. J Vis. 2019 Jun;19(6):13. doi:10.1167/19.6.13.

4. Ahuja A, Desrochers TM, Sheinberg DL. A role for visual areas in physics simulations. Cogn Neuropsychol. 2021 Oct-Dec;38(7-8):425–39. Epub 2022 Feb 13. doi:10.1080/02643294.2022.2034609.

5. Ahuja A, Yusif Rodriguez N, Ashok AK, Serre T, Desrochers TM, Sheinberg DL. Monkeys engage in visual simulation to solve complex problems. Curr Biol. 2024 Dec;34(24):5635-45.e3. doi:10.1016/j.cub.2024.10.026.

6. Rajalingham R, Sohn H, Jazayeri M. Dynamic tracking of objects in the macaque dorsomedial frontal cortex. Nat Commun. 2025 Jan;16(1):346. doi:10.1038/s41467-024-54688-y.

7. Wang Y, Ullman TD. Resource bounds on mental simulations: Evidence from a liquid-reasoning task. J Exp Psychol Gen. 2025 Aug;154(8):2105–24. doi:10.1037/xge0001792.

8. Ullman TD, Spelke E, Battaglia P, Tenenbaum JB. Mind games: Game engines as an architecture for intuitive physics. Trends in cognitive sciences. 2017;21(9):649–65.

9. Ludwin-Peery E, Bramley NR, Davis E, Gureckis TM. Limits on simulation approaches in intuitive physics. Cogn Psychol. 2021 Jun;127:101396. doi:10.1016/j.cogpsych.2021.101396.

10. Bass I, Smith KA, Bonawitz E, Ullman TD. Partial mental simulation explains fallacies in physical reasoning. Cogn Neuropsychol. 2021 Oct-Dec;38(7-8):413–24. Epub 2022 Jun 2. doi:10.1080/02643294.2022.2083950.

11. Li Y, Wang Y, Boger T, Smith KA, Gershman SJ, Ullman TD. An approximate representation of objects underlies physical reasoning. J Exp Psychol Gen. 2023 Nov;152(11):3074–86. doi:10.1037/xge0001439.

12. Sosa FA, Gershman SJ, Ullman TD. Blending simulation and abstraction for physical reasoning. Cognition. 2025 Jan;254:105995. doi:10.1016/j.cognition.2024.105995.

13. Smith KA, Battaglia PW, Tenenbaum JB. Integrating heuristic and simulation-based reasoning in intuitive physics [Preprint]. PsyArXiv; 2023 [cited 2026 Aug 18]. Available from: 10.31234/osf.io/bckes.

14. Griffiths TL, Lieder F, Goodman ND. Rational use of cognitive resources: Levels of analysis between the computational and the algorithmic. Top Cogn Sci. 2015 Apr;7(2):217–29. doi:10.1111/tops.12142.

15. Lieder F, Griffiths TL. Resource-rational analysis: Understanding human cognition as the optimal use of limited computational resources. Behav Brain Sci. 2020;43:e1. doi:10.1017/S0140525X1900061X.

16. Lieder F, Griffiths TL. Strategy selection as rational metareasoning. Psychol Rev. 2017 Nov;124(6):762–94. doi:10.1037/rev0000075.

17. Shenhav A, Botvinick MM, Cohen JD. The expected value of control: An integrative theory of anterior cingulate cortex function. Neuron. 2013 Jul;79(2):217–40. doi:10.1016/j.neuron.2013.07.007.

18. Callaway F, van Opheusden B, Gul S, Das P, Krueger PM, Griffiths TL, et al. Rational use of cognitive resources in human planning. Nat Hum Behav. 2022 Aug;6(8):1112–25. doi:10.1038/s41562-022-01332-8.

19. Ho MK, Abel D, Correa CG, Littman ML, Cohen JD, Griffiths TL. People construct simplified mental representations to plan. Nature. 2022 Jun;606(7912):129–36. doi:10.1038/s41586-022-04743-9.

20. Daw ND, Niv Y, Dayan P. Uncertainty-based competition between prefrontal and dorsolateral striatal systems for behavioral control. Nat Neurosci. 2005 Dec;8(12):1704–11. doi:10.1038/nn1560.

21. Lee SW, Shimojo S, O’Doherty JP. Neural computations underlying arbitration between model-based and model-free learning. Neuron. 2014 Feb;81(3):687–99. doi:10.1016/j.neuron.2013.11.028.

22. Kahn AE, Daw ND. Humans rationally balance detailed and temporally abstract world models. Commun Psychol. 2025 Jan;3(1):1. doi:10.1038/s44271-024-00169-3.

23. Kool W, Gershman SJ, Cushman FA. Cost-Benefit Arbitration Between Multiple Reinforcement-Learning Systems. Psychol Sci. 2017 Sep;28(9):1321–33. Epub 2017 Jul 21. doi:10.1177/0956797617708288.

24. Piloto LS, Weinstein A, Battaglia P, Botvinick M. Intuitive physics learning in a deep-learning model inspired by developmental psychology. Nat Hum Behav. 2022 Sep;6(9):1257–67. doi:10.1038/s41562-022-01394-8.

25. Nayebi A, Rajalingham R, Jazayeri M, Yang GR. Neural Foundations of Mental Simulation: Future Prediction of Latent Representations on Dynamic Scenes. In: Oh A, Naumann T, Globerson A, Saenko K, Hardt M, Levine S, editors. Advances in Neural Information Processing Systems. vol. 36. Curran Associates, Inc.; 2023. p. 70548–61. Available from: https://proceedings.neurips.cc/paper_files/paper/2023/file/df438caa36714f69277daa92d608dd63-Paper-Conference.pdf. doi:10.52202/075280-3091.

26. Ji-An L, Benna MK, Mattar MG. Discovering cognitive strategies with tiny recurrent neural networks. Nature. 2025 Aug;644(8078):993–1001. doi:10.1038/s41586-025-09142-4.

27. Gerstenberg T, Peterson MF, Goodman ND, Lagnado DA, Tenenbaum JB. Eye-tracking causality. Psychol Sci. 2017 Dec;28(12):1731–44. doi:10.1177/0956797617713053.

28. Krasich K, O’Neill K, De Brigard F. Looking at mental images: Eye-tracking mental simulation during retrospective causal judgment. Cogn Sci. 2024 Mar;48(3):e13426. doi:10.1111/cogs.13426.

29. van der Wel P, van Steenbergen H. Pupil dilation as an index of effort in cognitive control tasks: A review. Psychon Bull Rev. 2018 Dec;25(6):2005–15. doi:10.3758/s13423-018-1432-y.

30. da Silva Castanheira K, LoParco S, Otto AR. Task-evoked pupillary responses track effort exertion: Evidence from task-switching. Cogn Affect Behav Neurosci. 2021 Jun;21(3):592–606. doi:10.3758/s13415-020-00843-z.

31. Smith KA, Vul E. Sources of uncertainty in intuitive physics. Top Cogn Sci. 2013;5(1):185–99. doi:10.1111/tops.12009.

32. Pekkanen J, Lappi O. A New and General Approach to Signal Denoising and Eye Movement Classification Based on Segmented Linear Regression. Sci Rep. 2017;7:17726. doi:10.1038/s41598-017-17983-x.

33. Kowler E, Rubinstein JF, Santos EM, Wang J. Predictive smooth pursuit eye movements. Annu Rev Vis Sci. 2019 Sep;5:223–46. Epub 2019 Jul 5. doi:10.1146/annurev-vision-091718-014901.

34. Barnes GR. Cognitive processes involved in smooth pursuit eye movements. Brain Cogn. 2008;68(3):309–26. doi:10.1016/j.bandc.2008.08.020.

35. Joshi S, Gold JI. Pupil size as a window on neural substrates of cognition. Trends Cogn Sci. 2020 Jun;24(6):466–80. Epub 2020 Apr 21. doi:10.1016/j.tics.2020.03.005.

36. White CN, Poldrack RA. Using fMRI to constrain theories of cognition. Perspect Psychol Sci. 2013 Jan;8(1):79–83. doi:10.1177/1745691612469029.

37. Rieskamp J, Otto PE. SSL: A theory of how people learn to select strategies. J Exp Psychol Gen. 2006 May;135(2):207–36. doi:10.1037/0096-3445.135.2.207.

38. Callaway F, Gul S, Krueger PM, Griffiths TL, Lieder F. Learning to select computations. In: Proceedings of the 34th Conference on Uncertainty in Artificial Intelligence; 2018. p. 776–85. Available from: https://arxiv.org/abs/1711.06892.

39. Belledonne M, Butkus E, Scholl BJ, Yildirim I. Adaptive computation as a new mechanism of dynamic human attention. Psychol Rev. 2026;133(3):534–59. doi:10.1037/rev0000572.

40. Jensen KT, Hennequin G, Mattar MG. A recurrent network model of planning explains hippocampal replay and human behavior. Nat Neurosci. 2024 Jul;27(7):1340–8. doi:10.1038/s41593-024-01675-7.

41. Chen S, Callaway F, Kumar S, Lupkin SM, Wallis JD, McGinty VB, et al. Learning to select computations in recurrent neural circuits [Preprint]. bioRxiv; 2026 [cited 2026 Aug 18]. 2026.04.14.718499 [Version 1]. Available from: 10.64898/2026.04.14.718499.

42. Wang H, Jedoui K, Venkatesh R, Binder F, Tenenbaum JB, Fan JE, et al. Probabilistic simulation supports generalizable intuitive physics. In: Proceedings of the 46th Annual Meeting of the Cognitive Science Society; 2024. p. 1953–60. Available from: https://escholarship.org/uc/item/93j3f86q.

43. Mitko A, Navarro-Cebrián A, Cormiea S, Fischer J. A dedicated mental resource for intuitive physics. iScience. 2023 Dec;27(1):108607. doi:10.1016/j.isci.2023.108607.

44. Schwettmann S, Tenenbaum JB, Kanwisher N. Invariant representations of mass in the human brain. eLife. 2019;8:e46619. doi:10.7554/eLife.46619.

45. Pramod RT, Cohen MA, Tenenbaum JB, Kanwisher N. Invariant representation of physical stability in the human brain. eLife. 2022;11:e71736. doi:10.7554/eLife.71736.

46. Pramod RT, Mieczkowski E, Fang CX, Tenenbaum JB, Kanwisher N. Decoding predicted future states from the brain’s “physics engine”. Sci Adv. 2025 May;11(22):eadr7429. doi:10.1126/sciadv.adr7429.

47. Navarro-Cebrián A, Fischer J. Precise functional connections between the dorsal anterior cingulate cortex and areas recruited for physical inference. Eur J Neurosci. 2022;56(1):3660–73. doi:10.1111/ejn.15670.

48. Sosa FA, Zhang A, Gershman SJ, Ullman TD. Eye Gaze Reveals How People Flexibly Combine Abstraction and Mental Simulation [Preprint]. PsyArXiv; 2026 [cited 2026 Sep 2]. Available from: 10.31234/osf.io/6vyaz_v1.

49. Linsley D, Malik G, Kim J, Govindarajan LN, Mingolla E, Serre T. Tracking Without Re-recognition in Humans and Machines. In: Ranzato M, Beygelzimer A, Dauphin Y, Liang PS, Wortman Vaughan J, editors. Advances in Neural Information Processing Systems. vol. 34. Curran Associates, Inc.; 2021. p. 19473–86. Available from: https://proceedings.neurips.cc/paper/2021/hash/a2557a7b2e94197ff767970b67041697-Abstract.html.

